# Visual experience contributes to separation of face and language responses in the ventral stream

**DOI:** 10.1101/2024.12.18.629198

**Authors:** Elizabeth J. Saccone, Akshi, N. Apurva Ratan Murty, Judy S. Kim, Mengyu Tian, Nancy Kanwisher, Marina Bedny

**Author notes:** **Lead contact:** Elizabeth J. Saccone. **Corresponding author:** Marina Bedny.

## Abstract

Human ventral occipitotemporal cortex (vOTC) contains specialized regions that support visual recognition of behaviorally-relevant categories, including faces, written language and places (e.g., ^1–4^). An open question is how experience interacts with innate constraints to enable functional specialization. We investigate this question by comparing vOTC function across sighted and congenitally blind adults. In sighted adults, a region in lateral vOTC called the fusiform face area (FFA) responds preferentially to faces, whereas distinct left-lateralized portions of vOTC respond to written language ^1,2,5–12^. In blind people, lateral vOTC responds to face touching, braille and speech, but their functional co-localization has not been tested ^13–16^. The same group of congenitally blind adults (n=20) touched faces and spatial layouts (Experiment 1) and performed a reading (braille) and spoken language task (Experiment 2). Sighted adults performed analogous tasks in the visual modality (n=28). Using within subject analyses, we replicate the separation of faces and written language in sighted adults: written language responses are found only in left vOTC and within that hemisphere they are separate from faces. By contrast, left and right vOTC responds to language in people born blind and in the left hemisphere face and language responses overlap. These findings suggest that visual experience contributes to segregating responses to face and language in vOTC. Co-localization of face and language responses suggests an innate predisposition for communication-relevant processing in lateral vOTC.

## Results

In the first experiment, 20 blind participants touched 3D models of faces and scenes. In a second (language) experiment the same blind participants read braille words (tactile language) and felt shapes composed of braille dots (control). Sighted participants (n=15) viewed faces and objects (Experiment 1 sighted) and read (Experiment 2) printed words (visual language) and visual false fonts (control). Blind and sighted people also listened to spoken words (audio language) and backward speech (audio control) (Experiment 2; see Fig. 1).

**Fig 1.**
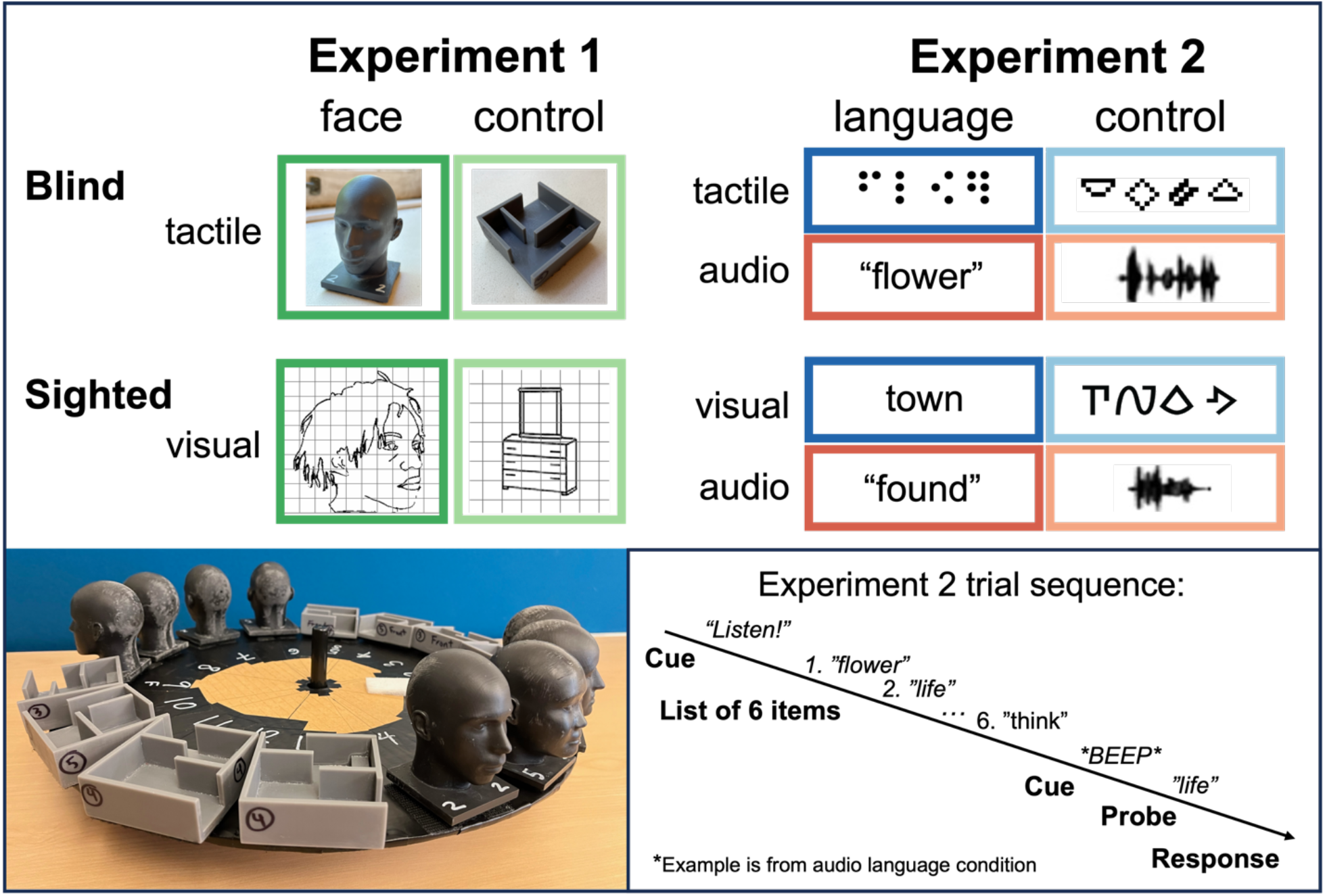
Experimental design and example stimuli. Control conditions were tactile scenes (Experiment 1 blind), visual objects (Experiment 1 sighted), tactile shapes (Experiment 2 blind), visual false fonts (Experiment 2 sighted) and audio backward speech (Experiment 2 blind and sighted).

### Lateral vOTC responds to tactile faces in people born blind (Experiment 1)

In an individual-subject functional region of interest (ROI) analysis (top 5% faces > control, leave-one-run-out cross validation), the lateral vOTC of both blind (tactile) and sighted (visual) participants preferred faces over control stimuli. We found a significant main effect of Condition and no Group-by-Condition interaction (Group (blind, sighted) x Hemisphere (left, right) x Condition (faces, control) mixed ANOVA in FFA parcel ^17^: main effect of Condition (faces vs. control), *F*(1,33)=55.3, *p*<.001; no Group-by-Condition interaction, *F*(1,33)=0.7, *p*=0.41; main effect of Group, *F*(1,33)=14.4, *p*<.001; see Supplementary Materials for all effects not reported in manuscript text) (Fig. 2, see Supplementary Fig. S1, S2 and S3 for results separated by hemisphere).

**Fig 2.**
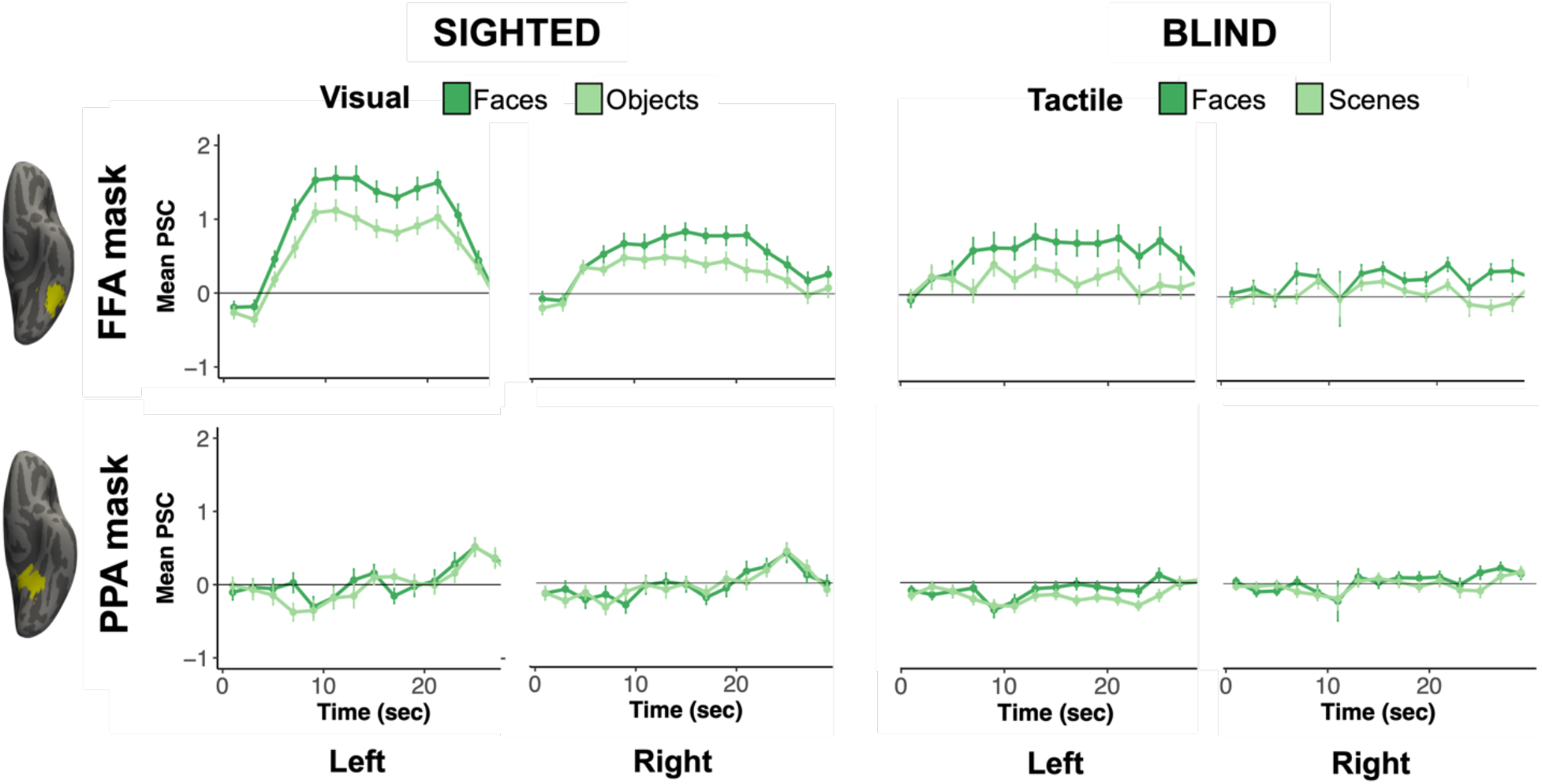
Responses to visual faces and objects (sighted) and tactile faces and scenes (blind) in face-preferring ROIs in left and right vOTC. Mean percent signal change (PSC) is plotted over time in face-preferring functional ROIs in FFA (top) and PPA (bottom) anatomical parcels from Julian et al. ^17^. ROIs were defined in individual participants using the top 5% of vertices for visual faces > objects (sighted) or tactile faces > scenes (blind) using a leave-one-run-out cross validation procedure. Error bars represent SEM.

Responses to faces were stronger than control stimuli for both blind and sighted groups independently (blind: Condition main effect (faces vs. scenes), *F*(1,19)=19.2, corrected *p*=.002; sighted: Condition main effect (faces vs. objects), *F*(1,14)=53.4, corrected *p*<.001).

There was also a marginal Hemisphere-by-Condition interaction (*F*(1,33)=3.6, *p*=.07) reflecting somewhat stronger responses to faces on the left. However, the three-way Group-by-Condition-by-Hemisphere interaction was not significant (*F*(1,33)=0.01, *p*=.9). Moreover, neither group showed a significant Condition-by-Hemisphere interaction independently. See Supplementary Materials for all within group ANOVA results.

Previous studies find that in sighted people faces activate lateral but not medial vOTC. We tested for this pattern in the blind group by comparing responses to tactile faces and scenes across lateral vOTC (visual face-responsive FFA) and medial vOTC (visual place-responsive parahippocampal place area; PPA ^3^) regions previously identified in sighted people ^17^. A Region (FFA parcel, PPA parcel) x Hemisphere (left, right) x Condition (face, control) repeated measures ANOVA revealed a Region-by-Condition interaction (*F*(1,18) = 19.1, *p*<.001), reflecting the fact that in people born blind, responses to faces are observed laterally but not medially in vOTC (Fig. 2). (A similar pattern was observed in the sighted control group (Fig. 2), see Supplemental Materials for details.) There were no interactions with Hemisphere (*p*s > .1).

In sum, blind individuals show a preference for tactile faces over tactile scenes in the canonical FFA location (lateral vOTC) but not the PPA location (medial vOTC), replicating earlier studies ^13^.

### Responses to faces and language overlap in vOTC of blind but not sighted (Experiment 2)

In sighted individuals, responses to faces and written language are observed in neighboring but separate portions of lateral vOTC ^8–11^. The so-called ‘visual word form area’ (VWFA) neighbors the FFA and responds preferentially to written language over matched objects and faces ^4,8,18–20^. In sighted people, responses to faces are typically larger in the right hemisphere, whereas responses to words are larger on the left. However, even within hemisphere, responses to faces and words are anatomically dissociable ^8,20^. Here we find that in people born blind, responses to faces and language are more overlapping in vOTC.

In face-preferring lateral vOTC, we observed a 3-way Group-by-Language condition-by-Modality interaction (Fig. 3) (*F*(1,33) = 7.8, *p*=.009), and a marginal 3-way Group-by-Hemisphere-by-Language condition interaction (*F*(1,33) = 4.1, *p*=.05), suggesting differences in face and language overlap across groups (mixed ANOVA in face-preferring ROIs in FFA parcel: Group (blind, sighted) x Hemisphere (left, right) x Language condition (words, control) x Modality (tactile/visual, audio)). We explore these interactions further below.

**Fig 3.**
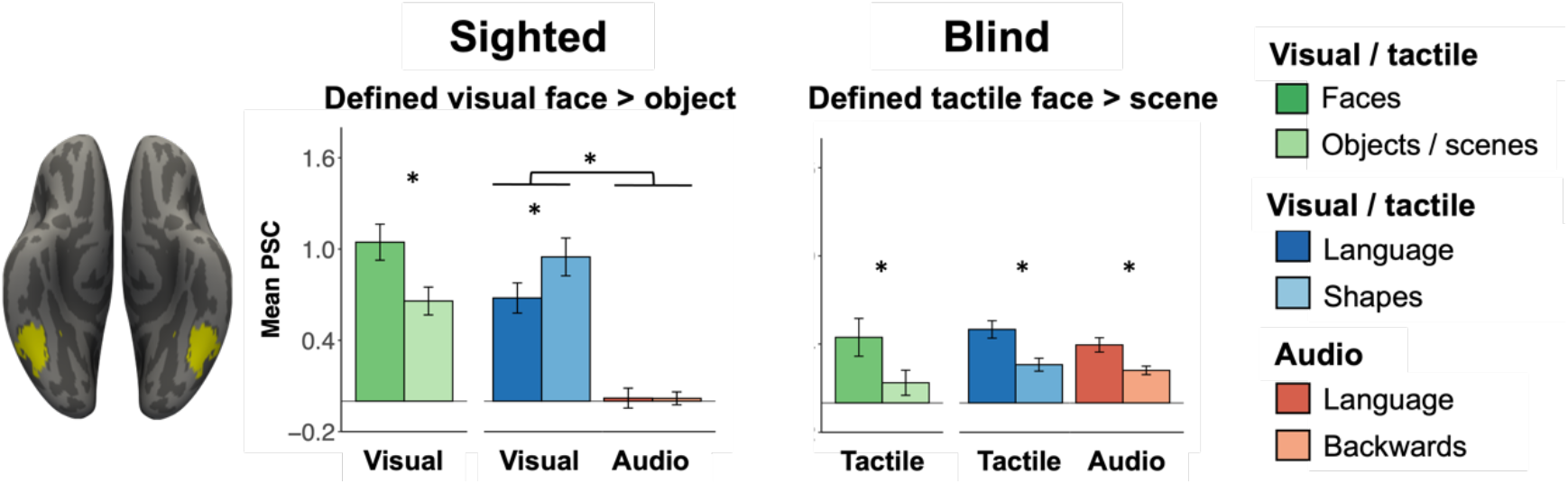
Responses to faces, written and audio words and control conditions in face-preferring functional ROIs in a FFA anatomical mask. Results include data from both hemispheres. Results separated by hemisphere are found in Fig. S1, S2 and S3 in the Supplementary Materials. Bars represent mean percent signal change (PSC) extracted for all conditions in ROIs defined within individual participants using the top 5% of vertices (k=12) for relevant contrast using a leave-one-run-out cross validation procedure. Error bars represent SEM. * *p*<.05 for post-hoc pairwise contrasts from within-group ROI analyses (see the main paper and Supplementary Materials) and for the main effect of Language condition in sighted face-preferring ROIs (top left). See Supplementary Materials for within-hemisphere comparisons of face and language effects (relative to control conditions) in face-preferring ROIs (Supplementary Results section 3).

### Sighted participants – within group ROI analysis

In sighted people, face-preferring bilateral FFA did not respond to either written or spoken language: indeed face-preferring vertices across both hemispheres responded more to false fonts than written language and showed no preference for spoken language over backward speech (main effect Language condition, *F*(1,14)=6.6, *p*=.02; visual false fonts > words, *t*(29)=3.6, corrected *p*=.002, Cohen’s *d*=0.4; audio language vs control, *t*(29)=0.03, corrected *p*>.99, Cohen’s *d*<0.01; Language condition-by-Modality interaction, *F*(1,14)=5.5, *p*=.04; no Hemisphere-by-Language condition-by-Modality interaction, *F*(1,14)=0.1, *p*=.75) (Fig. 3, Supplementary Fig. S1, S2). When an analysis was performed in each hemisphere separately comparing face and language effects (relative to control conditions), there were significant Experiment (face/language) by Condition (face/language vs. control) interactions in each hemisphere (Experiment-by-Condition interaction: left, *F*(1,14)=16.1, *p*=.001; right, *F*(1,14)=20.3, *p*<.001, see Supplementary Results section 3.1, Fig. S3 for details). For sighted people, face-preferring vertices do not respond to language in either hemisphere.

We next performed a control experiment (Experiment 3, see Supplementary Material for details) with a second group of sighted adults (n=15) where responses to visual faces were compared not only to objects but also to scenes (as well as bodies and scrambled objects) ^21^. The control experiment revealed highly similar selectivity profiles of the sighted FFA to that observed in Experiment 2. Face responsive vertices did not respond to language in either hemisphere, regardless of whether the FFA was defined as faces > scenes or faces > objects (top 5%, leave-one-run-out cross validation; see Fig. S4 in the Supplementary Materials).

Consistent with prior work ^8,20^, for sighted participants, written language-preferring vertices were found only in the left hemisphere (Figures S1, S3, S4). Written language-preferring vOTC vertices (ROIs defined visual words > false fonts in FFA parcel) also did not respond to faces over control objects in sighted people (Supplementary Fig. S1, S2). There was a marginal preference in the opposite direction: for objects over faces (main effect Condition, *F*(1,14)=3.7, *p*=.08; repeated measures ANOVA in written language-preferring ROIs in FFA parcel: Hemisphere (left, right) x Condition (faces, objects)). Selectivity for faces and language in vOTC of sighted people is anatomically separate in both hemispheres.

The sighted FFA also showed stronger responses to visual than auditory stimuli (main effect of Modality, *F*(1,14) = 37.4, *p*<.001; Hemisphere-by-Modality interaction, *F*(1,14)=9.3, *p*=.01; left hemisphere visual > audio, *t*(29)=7.8, corrected *p*>.001, Cohen’s *d* = 1.8; right hemisphere visual > audio, *t*(29)=5.4, corrected *p*>.001, Cohen’s *d*=1.1) (see Fig. 3). In sum, consistent with prior studies, in sighted people the FFA is a vision-dominated region that does not respond preferentially to written or spoken language.

### Blind participants – within group ROI analysis

In contrast, the face-preferring vOTC of blind participants responded to language (repeated measures ANOVA: Hemisphere (left, right) x Language condition (words, control) x Modality (tactile, audio), main effect Language condition *F*(1,19)=17.6, *p*<.001) (Fig. 3, Supplementary Fig. S1, S2). The effect of Language condition was qualified by a Hemisphere-by-Language condition interaction (*F*(1,19)=11.0, *p*=.004). The effect of language was significant in the left hemisphere face-preferring vertices (left hemisphere, Language condition effect *F*(1,19)=6.6, corrected *p*<.001) and marginal in the right hemisphere face-preferring vertices (right hemisphere, Language condition effect *F*(1,19)=2.1, corrected *p*=.09) (see Fig. 3, S1, S2).

Within-hemisphere comparisons of face responses (relative to control scenes) and language responses (relative to tactile shapes) in face-preferring vertices showed no significant interaction of Experiment (face/language) by Condition (face/language vs. control) in either hemisphere (main effect of Condition (faces/words > control condition): left, *F*(1,19)=23.6, *p*<.001; right, *F*(1,19)=19.6, *p*<.001), and no significant Experiment-by-Condition interactions: left, *F*(1,19)=0.1, *p*=.8; right, *F*(1,19)=0.8, *p*=.4, see Supplementary Fig. S3).

In other words, when face-preferring vertices are selected (faces > control), they show a response to language over control in blind individuals (see Fig. S3 and Supplementary Results section 3.2 for Experiment (1: face, 2: language) x Condition (key condition: faces/words, control condition: scenes/tactile shapes) repeated measures ANOVAs).

When we identified language-preferring vertices in bilateral FFA parcels (ROIs defined tactile language > shapes) we likewise observed a weak but reliable preference for faces over control scenes in the blind group (Supplementary Fig. S1, S2) (Condition main effect *F*(1,19)=6.0, *p*=.02 (repeated measures ANOVA: Hemisphere (left, right) x Condition (faces, scenes), no other effects were significant)). Inspection of the data in each hemisphere separately suggests that, for blind people, responses to faces are weaker in the right hemisphere (see Supplementary Fig. S1, S2).

All results were confirmed with a smaller smoothing kernel (2mm) and with ROIs defined instead using top 10% (k=24) vertices in the FFA anatomical parcel (see Fig. S5 in the Supplementary Materials).

In sum, sighted people show clear separation of face and language responses in both left and right vOTC. By contrast, there was more overlap between face and language responses in vOTC of blind people. People born blind showed responses to spoken and written language in both hemispheres and show overlap between faces and language within hemisphere. For blind people, the clearest within hemisphere overlap between faces and language is observed in the left hemisphere, where responses to faces and words were also stronger for the blind group.

### Responses to faces and language overlap in vOTC of blind group: Whole-cortex analysis

In whole-cortex analysis, left lateral vOTC of blind participants responded more to faces than scenes (cluster peak: −43, −46, −11, see Supplementary Table S2) (Fig. 4). This ‘FFA’ peak is broadly consistent with previously observed responses in sighted people, however, it is somewhat more anterior and left-lateralized (left FFA −40, −52, −18; right FFA 38, −42, − 22 ^17^). This pattern of slightly anterior left-lateralized peak in whole-cortex analysis is consistent with the whole-cortex participant-overlap map from a separate sample of blind participants ^13^. Although in ROI analyses, Ratan Murty et al. ^13^ observed effects in both hemispheres.

**Fig 4.**
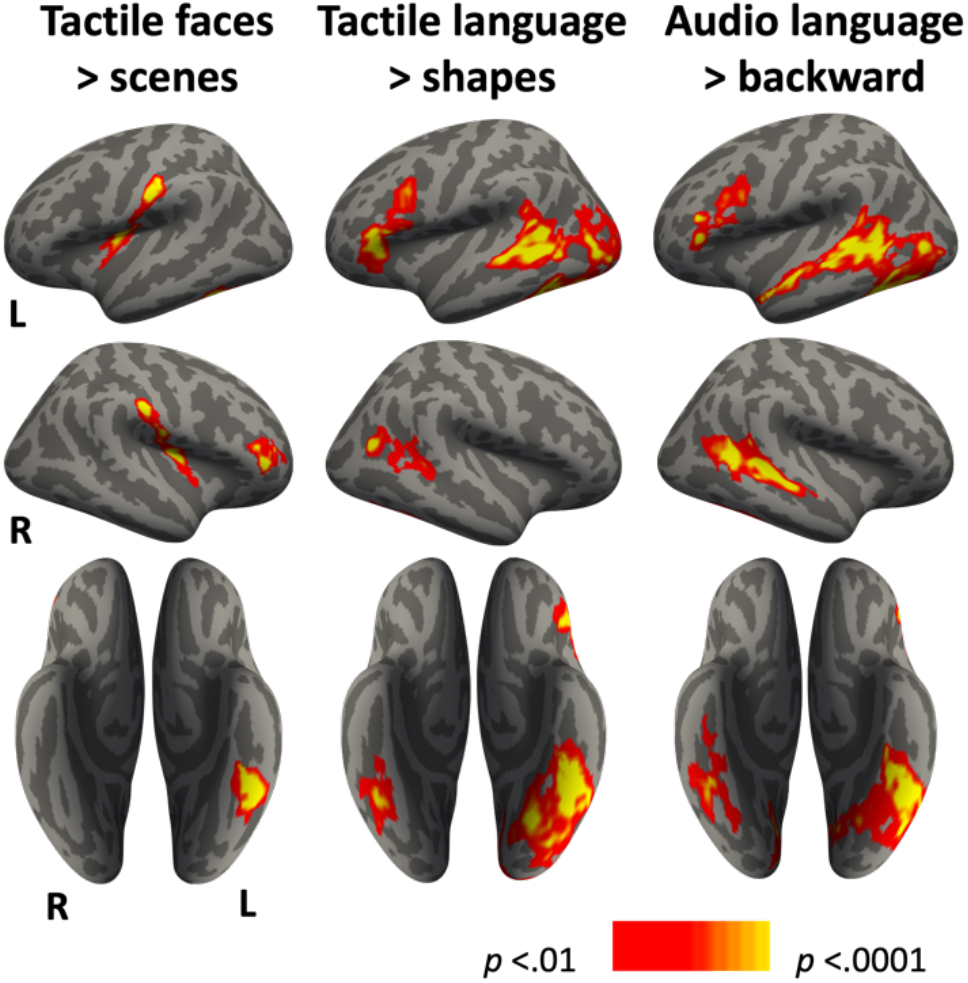
Whole-cortex analysis results in the blind group. BOLD responses for tactile faces > scenes (left column, Experiment 1), tactile language > control shapes (middle column, Experiment 2) and audio language > backward speech (right column, Experiment 2). Whole-brain random-effects analyses were run at the group level using mixed-effects models, thresholded *p<*0.05 cluster-wise and Family Wise Error corrected at *p<*0.01 vertex-wise for multiple comparisons using nonparametric permutation correction.

Left lateral vOTC of blind people also showed strong responses to tactile (braille) and spoken language (tactile language > shapes, audio language > backward speech). The language responses are in a similar location in the left hemisphere to the face responses but are stronger and more widespread, extending posteriorly into occipital regions and into the right lateral vOTC ^14–16,22–25^. The location of peak responses to faces and language in lateral vOTC was indistinguishable in an individual participant analysis comparing X, Y and Z coordinates in both hemispheres (all *p*s>.05, see Supplement for details).

In sum, in people born blind, the left lateral vOTC shows a distinctive functional profile, with a mixed selectivity for language and faces (Fig. 3, Supplementary Fig. S1, S2, S3). Outside of vOTC, both tactile (braille) and spoken language elicited robust activity in the canonical left frontotemporal language network of people born blind, extending posteriorly into occipital regions. By contrast, the strongest responses to touching faces were observed bilaterally in inferior parietal and postcentral regions, extending into the superior insulae (Fig. 4). In whole-cortex analysis, overlap between faces and language was observed in left vOTC of blind people but not elsewhere in the cortex.

## Discussion

Comparing lateral vOTC function across blind and sighted people reveals the interaction of innate constraints and experience during cortical development. We find more overlap between responses to faces and language in vOTC for blind than sighted people. Prior work finds that the lateral vOTC of literate sighted adults contains anatomically separable responses to faces and written language (e.g., ^8,12,20^). For sighted people, responses to written language are observed primarily in the left hemisphere, whereas responses to faces are bilateral and even within the left hemisphere, responses to face and written language are separate. We replicate this result in the current sample of sighted adults. By contrast, here we report that for people born blind, responses to written and spoken language are observed in both left and right vOTC and, in the left hemisphere, responses to faces and language overlap. In the right hemisphere, we did not observe as clear evidence for overlap in the blind group, but also failed to find clear evidence for separation of face and language responses, as we observed for the sighted. The current results suggest that vision facilitates the separation of face and language preferences in vOTC, both across and within hemispheres.

Unlike sighted people, people born blind do not use faces in any modality to recognize others and instead report relying on verbal and auditory cues. Indeed, blind people rate themselves as unlikely to be able to identify family members and close friends by touching their faces ^26^. One study also suggests that people who are blind are worse than sighted people at discriminating facial identity by touch, presumably because of lack of experience doing so ^27^. It is therefore all the more striking that lateral vOTC responds to faces over other tactile objects in this population, suggesting some form of vision-independent predisposition for faces in lateral vOTC. On the other hand, visual experience contributes to the separation of language as opposed to face specialization in vOTC.

One hypothesis is that lateral vOTC evolved as a visual component of a primate communication network. In sighted people, responses to faces and language in vOTC are anatomically proximal ^12,28–30^. Lateral vOTC shows connectivity to fronto-temporal language networks in infancy, long before literacy acquisition ^20,31^. Nonhuman primates rely heavily on vision during communication, using body and facial gestures, like lip-smacking ^32–34^. According to some views, human language evolved from such multi-modal communication ^35,36^. Consistent with this idea, although auditory languages are common, humans readily acquire visuo-manual, sign languages, which rely on both limb and facial gestures to signal lexical and grammatical information ^37–40^.

Connectivity between lateral vOTC and frontotemporal communication networks may enable this region to develop specialization for various communication-relevant visual functions, including recognition of faces and written language. In people born blind, this region becomes fully multi-modal and responds most strongly to spoken and written language. Together this evidence reveals how evolved anatomical constraints nevertheless leave room for cortical regions to adapt to the experience and behavioral needs of the individual.

## Conclusions

Comparison of vOTC function across sighted and blind adults reveals the interaction of innate constrains and experience during cortical specialization. Both sighted and blind adults show responses to faces in the lateral vOTC, implying vision-independent specialization. However, in people born blind there is more overlap between responses to faces and language, whereas in sighted people these responses are neighboring but distinct. Together with prior work, our findings point to an innate predisposition for communication-relevant processing in vOTC, with visual experience leading to further specialization for faces and language.

## Acknowledgments

The authors wish to thank the participants for their time, and the blind community and the National Federation for the Blind for their support. This work was supported by grants from Johns Hopkins Science of Learning Institute (80034917) and the NIH/NEI (R01 EY027352– 01).

## Author contributions

E.J.S.: Conceptualization, Methodology, Formal analysis, Investigation, Visualization, Project administration, Data curation, Writing – original draft, Writing - review and editing. A.: Conceptualization, Methodology, Software, Investigation, Writing - review and editing. N.A.R.M.: Conceptualization, Methodology, Software, Resources, Writing - review and editing. J.S.K.: Conceptualization, Methodology, Software, Investigation, Project administration, Writing - review and editing. M.T.: Formal analysis, Investigation, Project administration, Writing - review and editing. N.K.: Conceptualization, Methodology, Writing - review and editing. M.B.: Conceptualization, Resources, Supervision, Funding acquisition, Validation, Methodology, Project administration, Writing – original draft, Writing - review and editing.

## STAR Methods

**KEY RESOURCES TABLE**

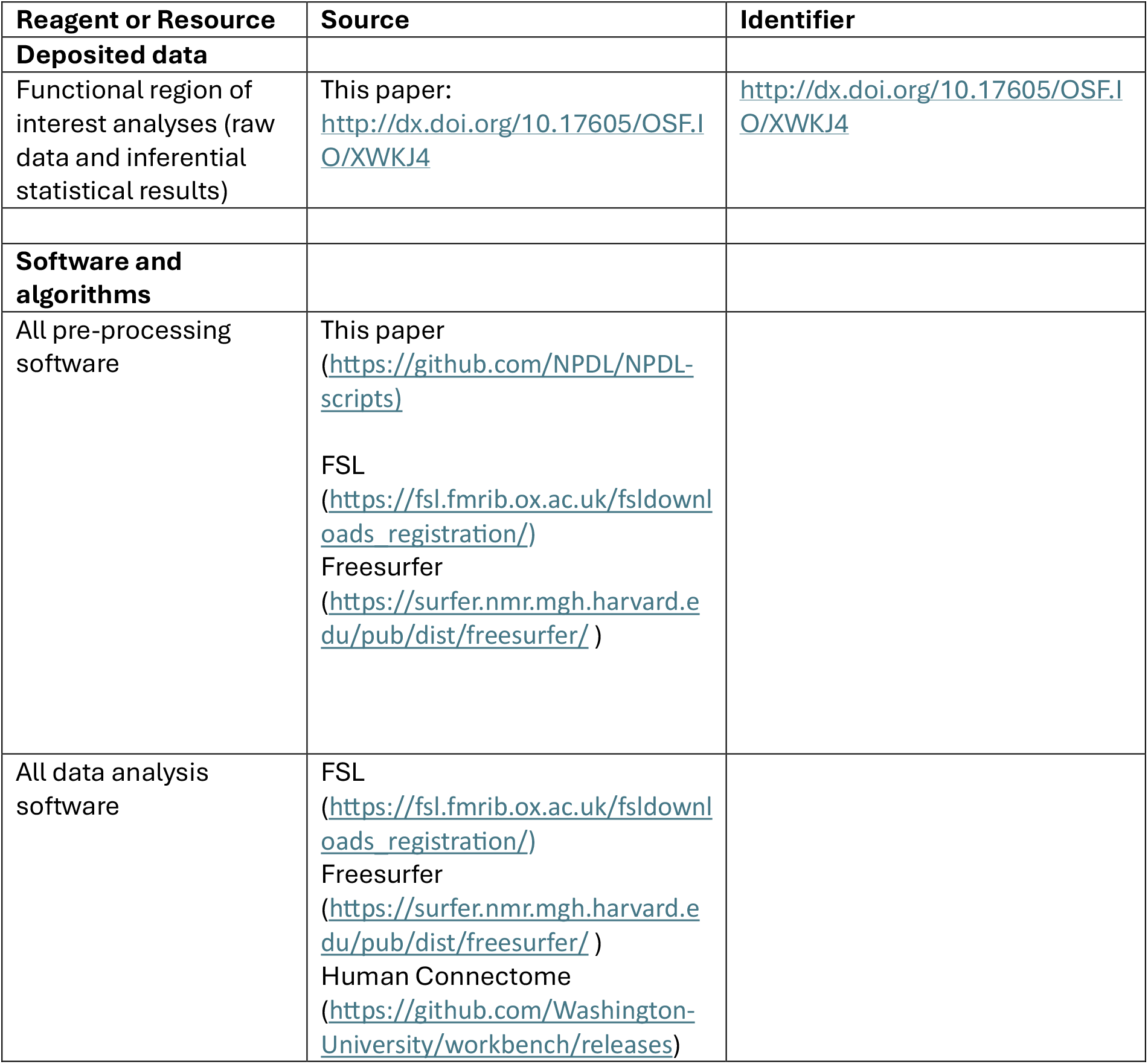
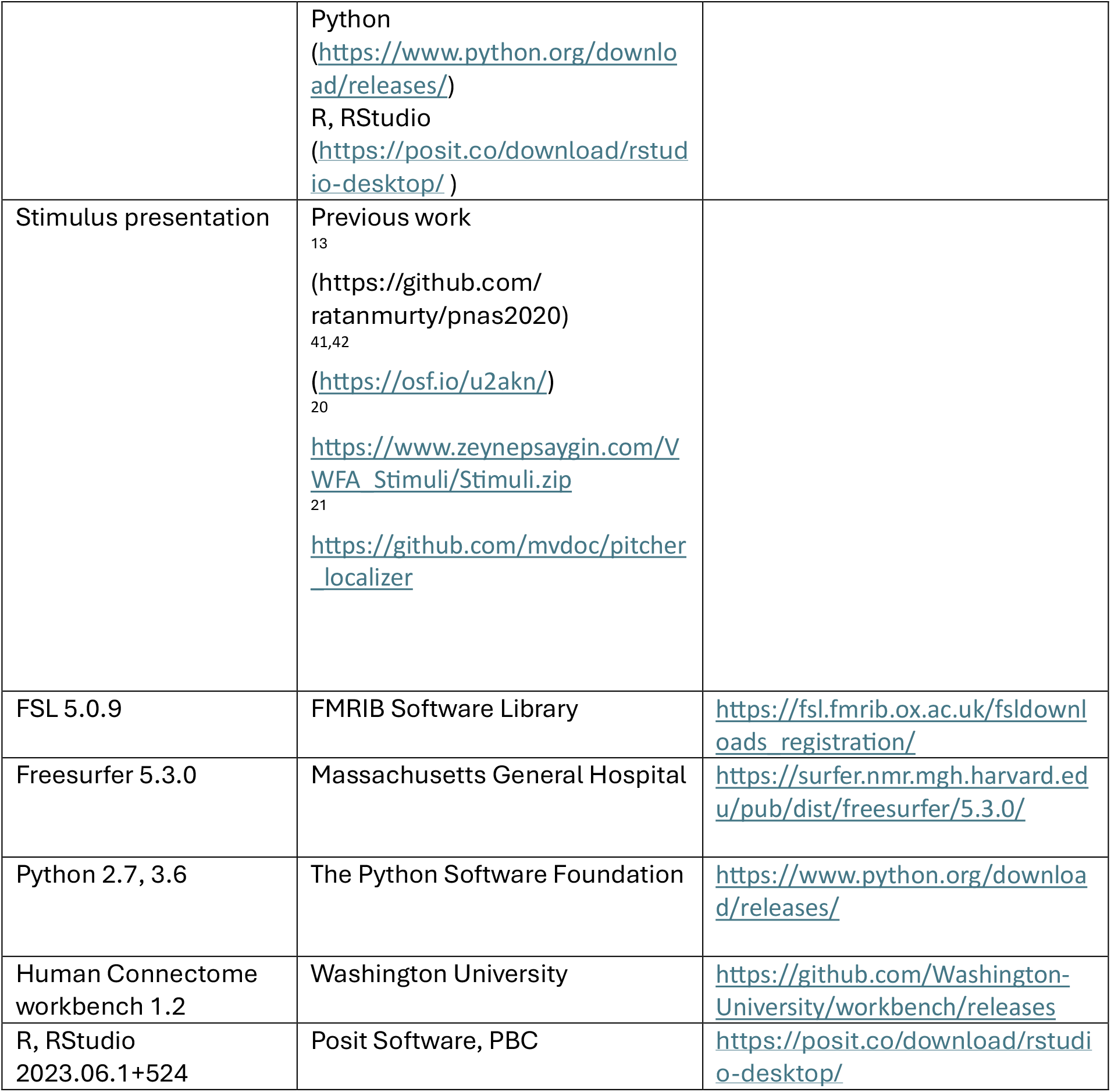

## RESOURCE AVAILABILITY

### Lead contact

Further information and requests for resources should be directed to and will be fulfilled by the lead contact, Elizabeth J. Saccone.

### Materials availability

This study did not generate new unique reagents.

## Data and code availability

All data and code on OSF, see Key Resources Table for doi

- The code and stimuli to run the tasks reported in this paper are available at the links reported in the Key Resources Table
- Extracted percent signal change (PSC) data from ROIs reported in this paper and code used to summarize and statistically analyze the PSC data are available at the project’s OSF page: http://dx.doi.org/10.17605/OSF.IO/XWKJ4
- The complete, standardized, de-identified fMRI data is available from the authors upon request, at a later date will be made publicly available

## EXPERIMENTAL MODEL AND SUBJECT DETAILS

Experiments 1 and 2 were completed by 20 congenitally blind adults (12 females, 8 males, mean age = 40.5 years, SD = 11.8 years) and 15 sighted adults (8 females, 6 males, 1 unspecified, mean age = 23.9 years, SD = 6.0 years) at Johns Hopkins’ F. M. Kirby Research Center in Baltimore, Maryland. Sighted adults were recruited from the Johns Hopkins University student and staff community and through public online advertisements. All participants were fluent English speakers, the majority of participants were native English speakers (90% of blind group; 73.3% (13.3% missing data) of sighted group). All participants were screened through self-report and all reported no cognitive or neurological disabilities. All blind participants were completely blind from birth, reporting at most minimal light perception. None reported having ever seen objects, shapes, colors or motion. All reported causes of blindness were pathology anterior to the optic chiasm i.e., were not due to cortical damage (see Supplementary Table S1). All had completed high school, with 18 (90%) having completed some college, 16 (80%) had completed a college degree. They were fluent braille readers who began learning braille on average at age 4.5 years (SD = 1.5 years) and rated their ability as mean 4.6 (SD = 0.8) from a scale from 1 (not at all well) to 5 (expert). All blind and sighted participants gave informed consent to participate according to procedures approved by the Johns Hopkins Institutional Review Board.

## METHOD DETAILS

### Stimuli and fMRI experimental procedures

#### Experiment 1 (blind group): Tactile face perception

Task and stimuli were taken from Ratan Murty et al. ^13^ except only face and scene conditions were included to reduce scan time (see Fig. 1). Participants touched one object at a time and performed 1-back repetition detection at the exemplar level. Objects were attached to a circular platform that the experimenter rotated every 6 seconds to present a new object to the participant’s hand, cued by an auditory tone delivered through headphones. Participants responded to every object, identifying it as a repeat or non-repeat. Half the participants touched the objects with their left hand and responded via button press with their right, and vice versa for the other half. Stimuli were presented in blocks of 4 objects from the same condition, creating 24-second blocks. Every block contained one repeated exemplar, with the identity repeated and the within-block position of the repeat counterbalanced across blocks and runs. There were 5 unique exemplars per condition.

Participants completed 5 runs following a short practice. Each run included 4 blocks per condition. For half of the participants, 3 runs began with faces and 2 began with scenes, with the reverse combination for the other half. There was a 50-second rest period in the middle of each run. For runs that began with a face block, the rest period was always followed by a scene block, and vice versa for runs that began with scenes. One participant only completed 3 runs due to time constraints.

#### Experiment 1 (sighted group): Visual face perception

The visual face localizer was taken from Saygin et al. ^20^. In brief, participants viewed line drawings of faces, objects (e.g., household objects, buildings), as well as words and scrambled words that were collected for a different study and not reported here. Participants performed 1-back repetition detection, with repeats indicated via button press. Stimuli from the same condition appeared in blocks of 28 items which included 2 repeated exemplars. Blocks were 18 seconds long. Participants completed 3 runs, each of which included 4 blocks per condition, as well as 3 rest blocks (fixation).

#### Experiment 2 (blind group): Language experiment

The language experiment was a delayed-match-to-sample paradigm described in detail elsewhere ^41,42^. In brief, trials comprised 6 stimuli from the same condition presented one at a time, followed by a beep, then a probe stimulus from the same condition (see Fig. 1). A response period followed where participants indicated via button press whether or not the probe stimulus had been in the initial list of six. Trials were either tactile or auditory for the blind group.

The tactile conditions were braille words, strings of 4 braille consonants (not reported here), or strings of 4 non-braille shapes that were comprised of braille pins (control). Braille words and characters were from Grade-II contracted braille, the most common form of braille in the United States. The braille words were on average 4 characters long (SD = 2.1), including contracted characters that represent more than one letter in visual print (e.g., -ed, -ing, the). Each tactile stimulus was presented for 2 seconds on a refreshable MRI-compatible tactile display. Three of the blind participants were from Kim et al.’s (2017) original study and had an interstimulus interval (ISI) of 0.75 seconds between stimuli, resulting in 6-item blocks that were 16.5 seconds. Due to a coding error, the rest of the blind participants performed the task with 1.75 seconds ISI, resulting in tactile blocks lasting 22.5 seconds (excluding the beep, probe stimulus and response period). Participants used one hand to perceive the tactile stimuli, based on their personal preference for braille reading, and responded with the other.

The auditory conditions were audio words or backward speech (words played in reverse: control) presented through MRI-compatible Sensimetrics headphones. Participants heard a different set of words than they read in the tactile word condition, but whether a particular word was read (tactile) versus heard (audio) was counterbalanced across participants. The auditory stimuli were 0.41 seconds long on average. Auditory blocks were approximately 16.5 seconds, designed to match the original tactile block length.

Each run included 4 blocks per condition as well as 6 × 16-second rest periods interspersed throughout the run. Condition order was counterbalanced across runs. There were 5 runs, leading to a total of 20 blocks per condition per participant. One participant only completed 4 runs due to time constraints.

#### Experiment 2 (sighted group): Language experiment

Sighted participants completed a comparable language experiment but with visual and auditory stimuli (see ^41,42^ for details). The visual conditions were American English words, strings of 4 consonants (not reported here), and strings of 4 non-letter shapes (false fonts: control). The words were on average 4 letters long (SD = 0.7). The non-letter control shapes were ‘false fonts’, designed to match English consonants on specific visual features ^41^. Visual stimuli were displayed centrally on a monitor, which the participants viewed via a mirror attached to the head coil. Because visual reading is considerably faster than tactile reading, the visual stimuli were presented for 1 second each with an ISI of 0.75, resulting in visual condition blocks of 10.5 seconds. The auditory stimuli were comparable to those described for the blind group but were presented with shorter ISI to match the visual block lengths of 10.5 seconds. One sighted participant only completed 4 of the 5 runs.

### fMRI data acquisition

All images were acquired using a 3T Phillips scanner. T1-weighted images were collected using a magnetization-prepared rapid gradient-echo (MP-RAGE) in 150 axial slices with 1mm isotropic voxels. Functional BOLD scans were collected in 36 sequential ascending axial slices. TR = 2s, TE = 30ms, flip angle = 70°, voxel size = 2.4 × 2.4 × 2.5mm, inter-slice gap = 0.5mm, field of view = 192 × 172.8 × 107.5.

### fMRI data analysis

#### Preprocessing

Analyses were performed using FSL (version5.0.9), FreeSurfer (version5.3.0), the Human Connectome Project workbench (version1.2.0), and custom in-house software (https://github.com/NPDL/NPDL-scripts). The cortical surface was created for each participant using the standard FreeSurfer pipeline ^43–45^. For task data, preprocessing of functional data included motion-correction, high-pass filtering (128s cut-off), and resampling to the cortical surface. Cerebellar and subcortical structures were excluded. The task data were smoothed on the surface with an 8.24 mm FWHM Gaussian kernel. Key analyses were repeated with a 2mm smoothing kernel and produced the same results (see Supplementary Fig. S5).

For each experiment, conditions were included as covariates of interest in general linear models, convolved with a standard hemodynamic response function (HRF) including temporal derivatives. Conditions for the blind participants were tactile faces and tactile scenes (Experiment 1) and tactile words, tactile shapes, audio words and backward speech (Experiment 2). Conditions for the sighted participants were visual faces and visual objects (Experiment 1) and visual words, visual false fonts, audio words and backward speech (Experiment 2). Data were pre-whitened to remove temporal autocorrelation. For Experiment 2 (language experiments), response periods and trials where participants did not respond were modeled with separate regressors. White matter signal, CSF signal, and timepoints with excessive motion (FDRMS > 1.5mm) were also included as covariates of no interest. One blind participant moved excessively during most runs and so an aggressive strategy was taken to remove motion-affected time points for this individual (162 timepoints were removed). After this process, visual inspection of this individual’s data showed activation patterns consistent with a canonical HRF, so their data were included. Aside from this participant, mean number of timepoints removed per participant due to motion were 2.1 (SD=3.2) for the blind group and 0.9 (SD=1.9) for the sighted group. Runs were combined within subjects using fixed-effects models.

#### Functional ROI analyses

We defined functional individual-subject regions of interest (ROIs) in the lateral vOTC canonical fusiform face area (FFA) location and in the medial vOTC canonical parahippocampal place area (PPA) location. Search spaces for both the FFA and the PPA ROIs were taken from a study of 35 sighted individuals looking at faces and scenes versus control conditions ^17^ (https://web.mit.edu/bcs/nklab/GSS.shtml). For the FFA search space, given that the left hemisphere FFA parcel was quite small, we flipped the right hemisphere parcel for all left hemisphere analyses (see ^8^ for a similar approach).

We defined face-preferring ROIs in individual participants with the faces > control contrasts in both groups (blind: tactile faces > scenes; sighted: visual faces > objects). The language preferring ROIs were defined using the written words > control shapes contrast (blind: tactile words > shapes; sighted: visual words > false fonts).

Each ROI was defined as the top 5% of vertices activated for each contrast within the particular anatomical mask, with signal then extracted using a leave-one-run-out cross validation procedure to ensure statistical independence. Specifically, ROIs were defined for a particular contrast (e.g., tactile face-preferring ROIs were defined tactile faces > scenes) using all but one run and this was repeated iteratively for all combinations of runs. Then the percent signal change (PSC: *Signal condition – Signal rest)/Signal rest*) was extracted from the left-out run and averaged for every condition, within and across runs. This process allows us to identify the most relevant signal for a contrast of interest at an individual level, while also ensuring that independent data are used to define and test ROIs ^46,47^. For all conditions we averaged PSC over the entire task block, after dropping the first 3 time points of the block to allow for the hemodynamic signal to rise.

Mixed-effects ANOVAs tested for group and condition differences. Post-hoc t-tests were performed and Bonferroni corrected *p*-values are reported where necessary to correct for multiple comparisons ensuring a family-wise *p*-value error rate of *p*<.05. Cohen’s d effect sizes are reported for post-hoc *t*-tests.

For the tactile face-preferring ROIs, there was one case of an outlier. Specifically, one blind participant had very high (> 3SD) PSC values for the tactile face and scene conditions in their most face-preferring vertices in the left hemisphere FFA mask location. Visual inspection of the data confirmed an appropriate hemodynamic response pattern, but with very high PSC values for this ROI only. Their data are included in the manuscript, although the pattern and statistical significance of the key findings are identical with the outlier data excluded. For a different blind participant, we could not define tactile face-preferring ROIs in the PPA parcel.

#### Whole-brain analyses

Whole-brain random-effects analyses were run at the group level using mixed-effects models, thresholded *p<*0.05 cluster-wise and Family Wise Error corrected at *p<*0.01 vertex-wise for multiple comparisons using nonparametric permutation correction ^48–50^. Whole-brain cluster peak locations can be found in Table S2 in the Supplementary Materials.

#### Individual peak comparison analysis

The MNI152 X, Y and Z coordinates of each individual’s peak responses to faces, braille words and audio words relative to control conditions were identified and compared to test for differences in their spatial distribution using separate one-way ANOVAs. Within a vOTC mask we identified the vertex location of the peak z-value for the three key contrasts (tactile faces > scenes, tactile word > shapes and audio word > backward speech). Following prior studies, a vOTC mask was defined by combining FreeSurfer atlas-based labels for the fusiform gyrus, parahippocampal and infero-temporal cortices ^51,52^.

