## Supplementary Material for "Visual experience contributes to separation of face and language responses in the ventral stream"

**Supplementary Figures**

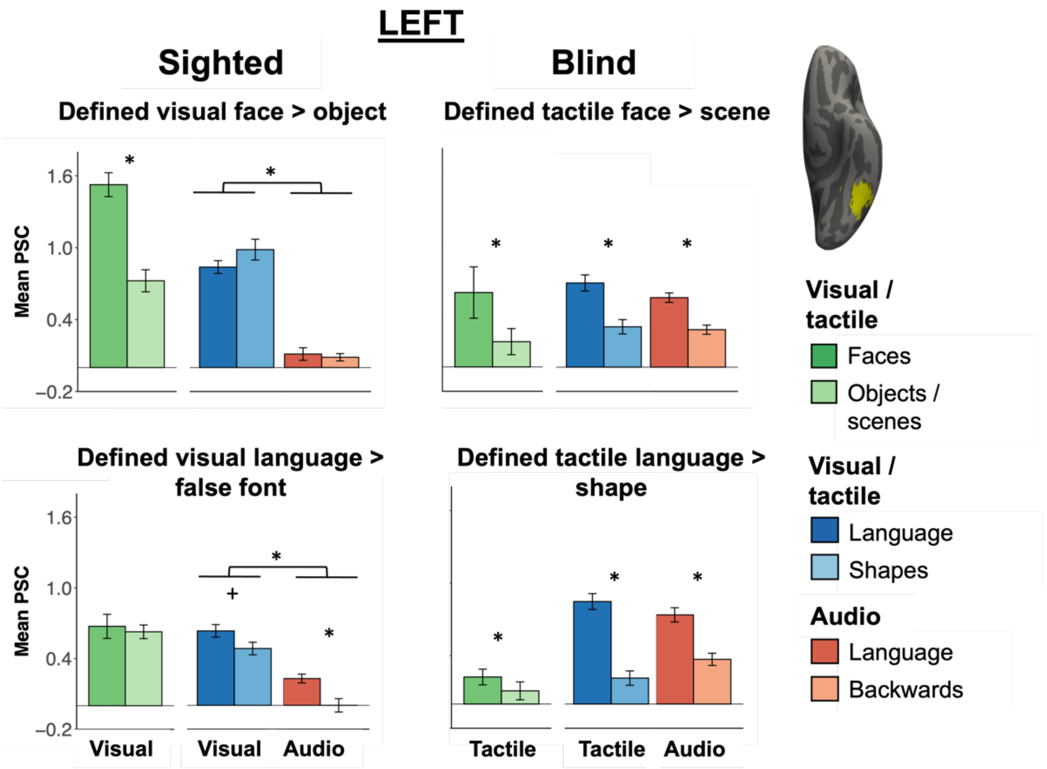


**Fig. S1.** Responses to faces, written and audio words and control conditions in face- (top) and language-preferring (bottom) functional ROIs in a left hemisphere FFA anatomical mask. Here, the sighted data are pooled from both sighted groups (n=28 in total completed Experiments 1 or 3 and also Experiment 2 (2 participants completed both Experiments 1 and 3 so only their data from Experiment 1 were included in this figure)). Control conditions for the blind group were tactile scenes, tactile shapes and audio backward speech. Condition conditions for the sighted group were visual objects, visual false fonts and audio backward speech. Bars represent mean percent signal change (PSC) extracted for all conditions in ROIs defined within individual participants using the top 5% of vertices (k=12) for relevant contrast using a leave-one-run-out cross validation procedure. Error bars represent SEM. * denotes significance at p<.05 and + denotes marginal significance (p =.09) for main effect of Modality (sighted group only) or post-hoc pairwise contrasts from within group ROI analyses (see main paper and Supplementary Results sections 2.2 and 2.3).

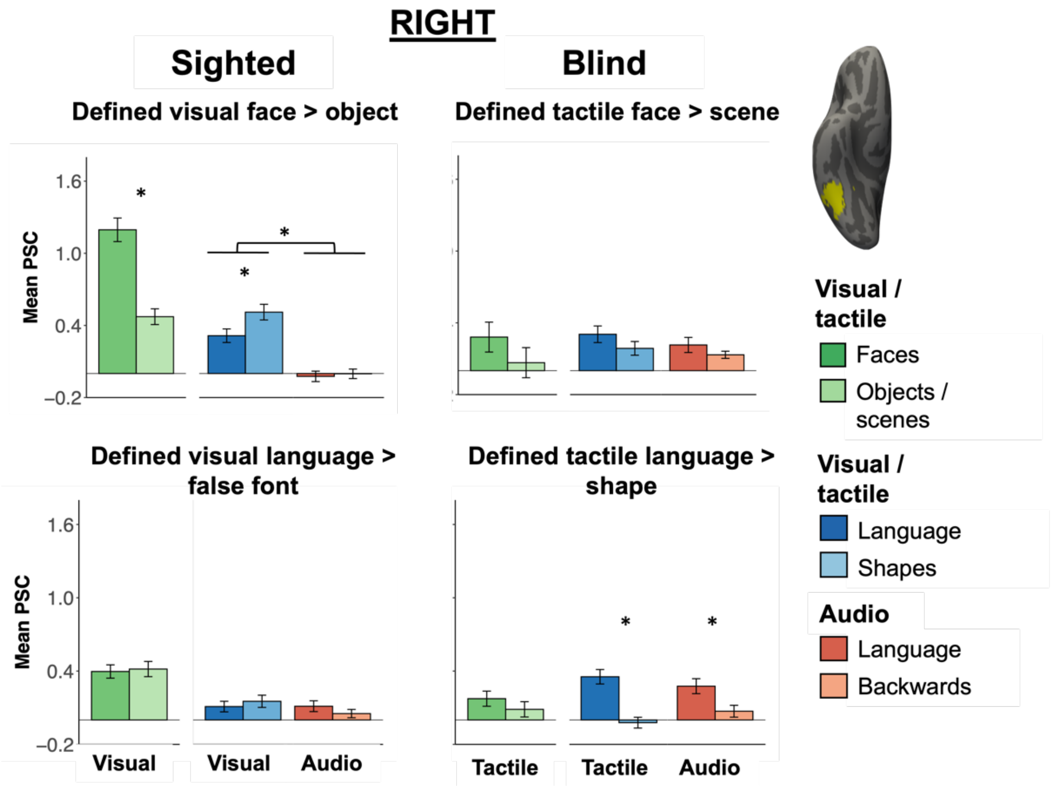


**Fig. S2.** Responses to faces, written and audio words and control conditions in face- (top) and language-preferring (bottom) functional ROIs in a right hemisphere FFA anatomical mask. Here, the sighted data are pooled from both sighted groups (n=28 in total completed Experiments 1 or 3 and also Experiment 2 (2 participants completed both Experiments 1 and 3 so only their data from Experiment 1 were included in this figure)). Control conditions for the blind group were tactile scenes, tactile shapes and audio backward speech. Condition conditions for the sighted group were visual objects, visual false fonts and audio backward speech. Bars represent mean percent signal change (PSC) extracted for all conditions in ROIs defined within individual participants using the top 5% of vertices (k=12) for relevant contrast using a leave-one-run-out cross validation procedure. Error bars represent SEM. * denotes significance at p<.05 for main effect of Modality (sighted group only) or post-hoc pairwise contrasts from within group ROI analyses (see main paper and Supplementary Results sections 2.2 and 2.3).

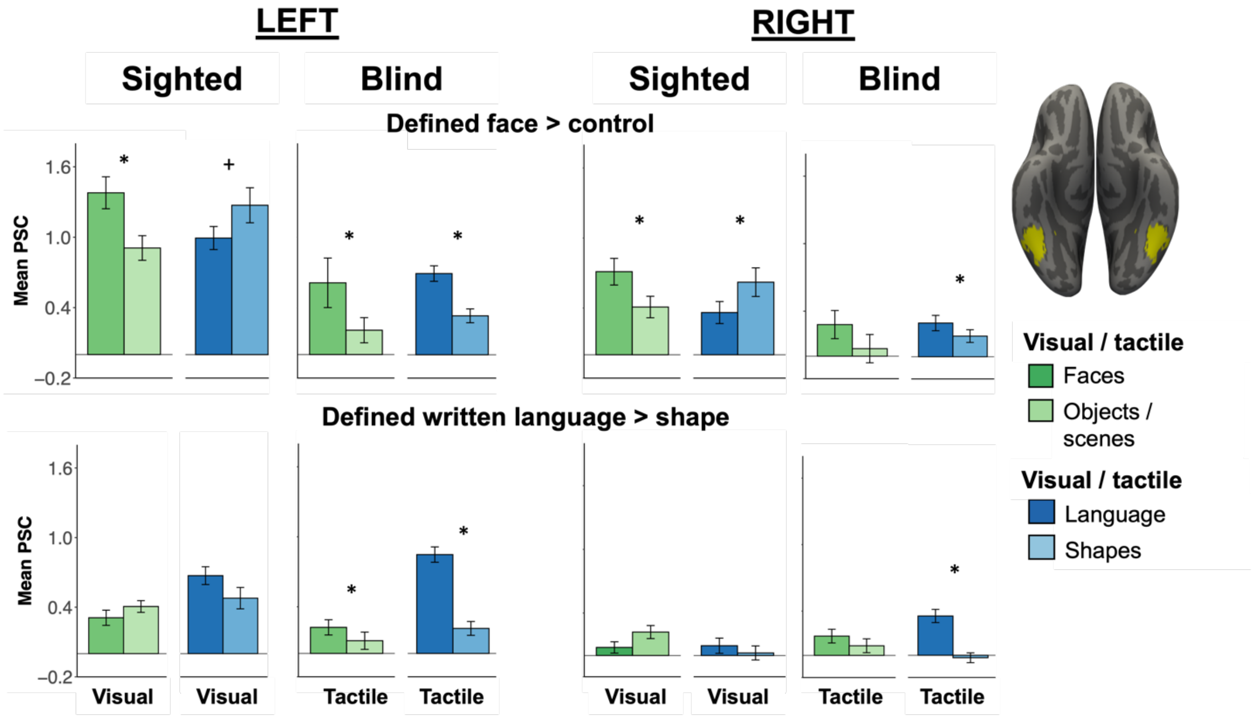


**Figure S3.** Responses to faces and written (visual or tactile) language and control conditions (Experiments 1 and 2) in face- (top) and visual/tactile language-preferring (bottom) functional ROIs in left and right FFA anatomical mask. Bars represent mean percent signal change (PSC) extracted for all conditions in ROIs defined within individual participants using the top 5% of vertices (k=12) for relevant contrast using a leave-one-run-out cross validation procedure. Error bars represent SEM. * denotes significance at $p<.05$ and + denotes marginal significance ($p = .09$) for post-hoc pairwise comparisons from Experiment (1: face experiment; 2: language experiment) x Condition (key condition: faces/words; control condition: objects or scenes/shapes) repeated measures ANOVAs performed within hemispheres (see Supplementary Results sections 3.1 (sighted) and 3.2 (blind) for all results).

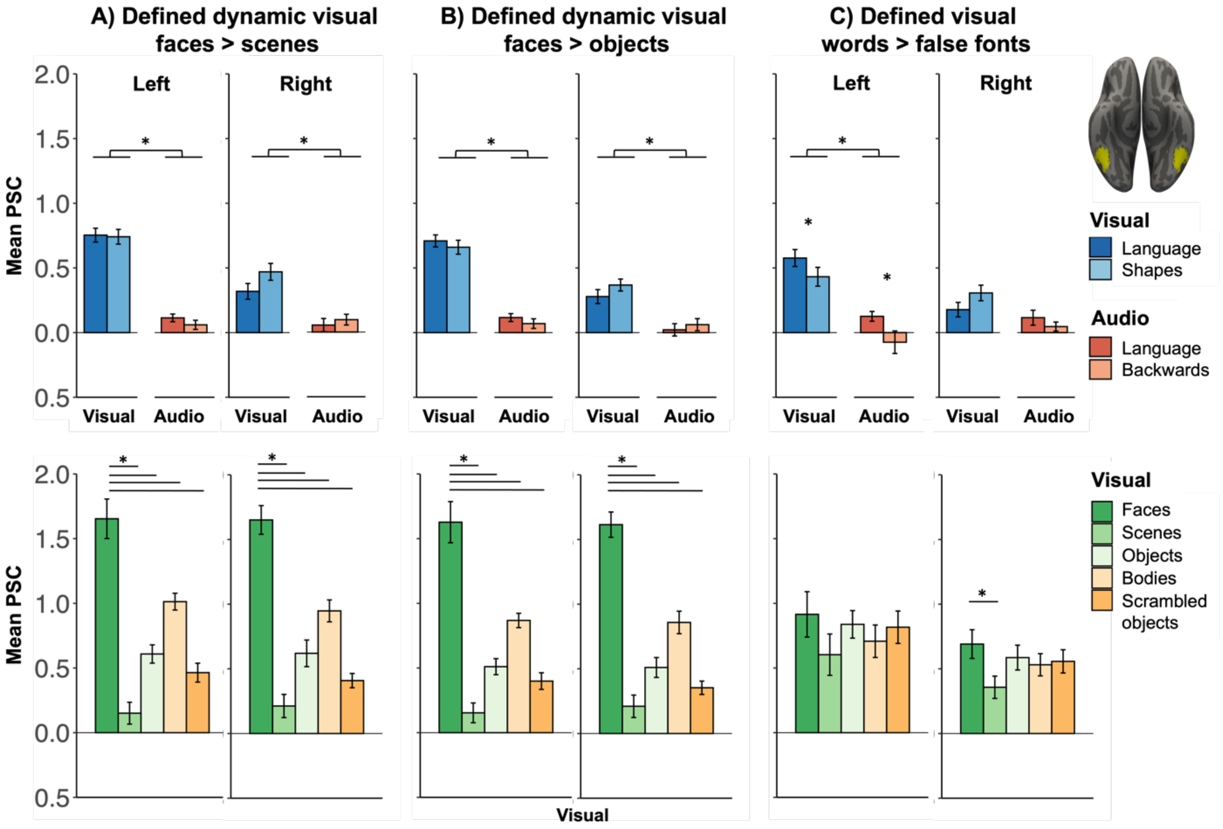


**Figure S4.** Separate responses to language (top) and faces (bottom) in Experiment 3 (additional group of sighted adults; n=15; age (years) mean = 41.7, SD=14.2; education (years) mean = 17.2 y, SD=2.3) in functional ROIs in anatomical FFA parcel. Experimental procedures and analysis methods are the same as that reported in the main paper, except that the face viewing task was a dynamic face localizer from Pitcher et al. (2011). In brief, participants performed 1-back repetition detection with short video clips of faces, bodies, objects, scenes and scrambled objects. ROIs were defined using the top 5% of **(A)** faces > scene contrast, **(B)** faces > objects contrast, and **(C)** visual words > false fonts from the same task described in Experiment 2. Bars represent mean percent signal change (PSC) extracted for all conditions using independent data (leave-one-run-out cross validation). Error bars represent SEM. See Supplementary Results section 4 for detailed results. Top row: * denotes significance at p<.05 for post-hoc pairwise comparisons and main effects of Language condition from Language condition (language, control) by Modality (visual, audio) repeated measures ANOVAs performed within hemispheres (see Supplementary Results section 4.2 for all effects). Bottom row: * denotes significance at p<.05 post-hoc pairwise comparisons between faces and other conditions where there was an effect of Condition (faces, scenes, objects, bodies, scrambled objects) in one-way ANOVAs performed within hemispheres (see Supplementary Results section 4.1).

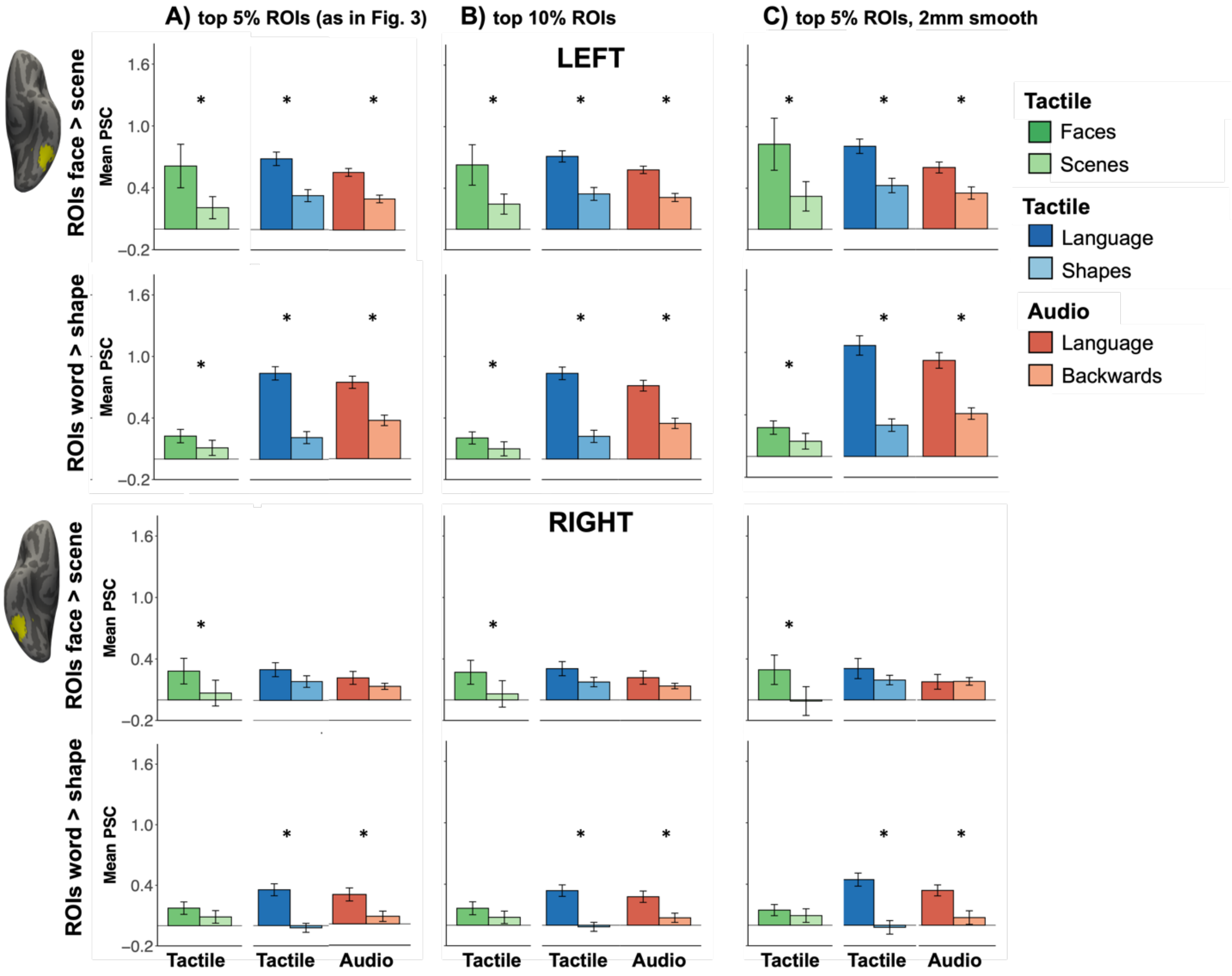


**Figure S5.** Blind group's responses to faces, written and audio words and control conditions in face- (rows 1 and 3) and tactile word-preferring (rows 2 and 4) functional ROIs in a FFA anatomical mask (Experiments 1 and 2) (Julian et al., 2012). Bars represent mean percent signal change (PSC) extracted for all conditions in ROIs defined within individual participants using a leave-one-run-out cross validation procedure. ROIs were defined using: **(A)** the top 5% of vertices (k=12) for the relevant contrast (as reported in the main manuscript and in Fig. S1, S2; see Fig. 3 for results collapsed across hemisphere), **(B)** the top 10% of vertices (k=24) for the relevant contrast, and **(C)** using the top 5% of vertices (k=12) for the relevant contrast and data smoothed on the surface with a smaller smoothing kernel (2mm). Error bars represent SEM. * denotes significance at $p<.05$ for paired samples t-tests for the face effect (tactile faces vs tactile scenes from Experiment 1 (green bars) or for post-hoc pairwise comparisons from Language condition (language, control) by Modality (tactile, audio) repeated measures ANOVAs from Experiment 2 (blue and pink bars).

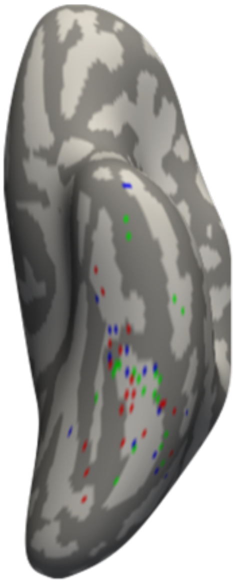

**Fig. S6.** Left hemisphere vOTC locations of blind participants' peak contrast values for tactile faces > scenes (green), tactile language > shapes (blue) and audio language > backward speech (pink) (Experiments 1 and 2). Each value is based on individual participants' fixed-effects results.

### Supplementary Results

*1.1 Lateral vOTC responds to tactile faces in people born blind (Experiment 1)*

In an individual-subject functional region of interest (ROI) analysis (top 5% faces > control, leave-one-run-out cross validation), the lateral vOTC of both blind (tactile) and sighted (visual) participants preferred faces over control stimuli (Group (blind, sighted) x Hemisphere (left, right) x Condition (faces, control) mixed ANOVA in FFA parcel). There were significant main effects of Group (F(1,33)=14.4, p <.001), Hemisphere (F(1,33)=5.5, p=.03), and Condition (F(1,33)=55.3, p<.001), reflecting stronger overall responses in the sighted group, in the left hemisphere, and for faces rather than control stimuli. None of the interactions were significant (Group-by-Condition, F(1,33)=0.7, p=.41; Group-by-Hemisphere, F(1,33)=1.0, p=.33; Condition-by-Hemisphere, F(1,33)=3.6,p=.07; Group-by-Condition-by-Hemisphere, F(1,33)<0.01, p=.91).

Although there were no interactions with Group, for completeness a Hemisphere (left, right) x Condition (faces, control) repeated measures ANOVA was run for each group separately. For the sighted group, there were main effects of Hemisphere (F(1,14)=9.0, corrected p=.02) and Condition (F(1,14)=53.4, corrected p<.001), once again reflecting stronger overall responses in the left hemisphere and to faces rather than control objects. The Hemisphere-by-Condition interaction was not significant (F(1,14)=2.4, corrected p=.28).

For the blind group, there was a significant main effect of Condition (F(1,19)=19.2, corrected p=.002), reflecting stronger responses to tactile faces than scenes. There was no significant main effect of Hemisphere (F(1,19)=0.8, corrected p=.72) and no Hemisphere-by-Condition interaction (F(1,19)=1.8, corrected p=.38).

*1.2 Face responses are found in lateral but not medial vOTC in both blind and sighted groups*

Next, for the blind group, we tested if there were stronger tactile face-preferring responses in lateral vOTC (FFA location) than in medial vOTC (PPA location), consistent with the typical pattern in the sighted brain. A Region (FFA parcel, PPA parcel) x Hemisphere (left, right) x Condition (faces, scenes) repeated measures ANOVA was performed on face-responsive vertices (top 5% faces > control, leave-one-run-out cross validation). Note that a face-preferring ROI in the right PPA parcel could not be defined for one blind participant. There was significant main effect of Region (F(1,18)=30.5, p<.001), reflecting stronger responses overall in the (lateral) FFA parcel than in the (medial) PPA parcel. The main effect of Condition was also significant (F(1,18)=19.1, p<.001), with stronger responses to tactile faces than scenes. The main effect of Hemisphere was not significant (F(1,18)=0.2, p=.68). There was a significant Region-by-Condition interaction (F(1,18)=19.1, p<.001) – but there was a significant difference between faces and scenes in both the FFA parcel (t(40)=4.3, corrected p<.001) and the PPA parcel (t(39)=3.2, corrected p=.01). Only in the FFA parcel were responses to faces above rest (one-sample t-test in FFA parcel: t(39)=3.2, p=.002; one-sample t-test in PPA parcel: t(38)=0.6, p=.7). None of the other interactions were significant (Region-by-Hemisphere, F(1,18)=3.3, p=.09; Hemisphere-by-Condition, F(1,18)=2.6, p.13; Region-by-Hemisphere-by-Condition, F(1,18)=0.8, p=.37).

As a control analysis, we repeated the above test for the sighted group, which confirmed the typical pattern of face-preferring responses in lateral but not medial vOTC (Region (FFA parcel, PPA parcel) x Hemisphere (left, right) x Condition (faces, objects) repeated measures ANOVA). Note that a face-preferring ROI in the left PPA parcel could not be defined in one sighted participant. There were significant main effects of Region (F(1,13)=79.0, p<.001), Hemisphere (F(1,13)=8.8, p=.01), and Condition (F(1,13)=17.4, p=.001), reflecting stronger responses overall in the (lateral) FFA parcel than in the (medial) PPA parcel, in the left hemisphere, and for faces over control objects. There was a significant Region-by-Condition interaction (F(1,13)=24.3, p<.001), with stronger responses to faces over objects in the FFA parcel (t(30)=7.1, corrected p<.001) but not in the PPA parcel (t(29)=0.4, corrected p>.99). The Region-by-Hemisphere interaction was marginal (F(1,13)=4.6, p=.05). None of the other interactions were significant (Hemisphere-by-Condition F(1,13)=2.4, p=.15; Region-by-Hemisphere-by-Condition F(1,13)=0.1, p=.8).

*2.1 Responses to faces and language overlap in vOTC of blind but not sighted (Experiment 2)*

We next tested responses to the language conditions in the face-preferring ROIs in FFA parcels using a Group (blind, sighted) x Hemisphere (left, right) x Language condition (words, control) x Modality (tactile/visual, audio) mixed ANOVA. There was a significant main effect of Hemisphere (F(1,33)=25.2, p<.001), with stronger responses in the left hemisphere (see Fig. S1 and S2). The main effect of Modality was also significant (F(1,33)=45.3, p<.001), with stronger responses to visual/tactile stimuli than auditory. There were no significant main effects of Group (F(1,33)=0.6, p=.44) or Language condition (F(1,33)=1.0, p=.33).

A number of two-way interactions were significant (Hemisphere-by-Modality interaction F(1,33)=8.9, p=.005; Hemisphere-by-Language condition interaction F(1,33)=4.7, p=.04; Group-by-Language condition interaction F(1,33)=22.0, p<.001; Group-by-Modality interaction F(1,33)=31.5, p<.001). There were significant three-way interactions of Group-by-Hemisphere-by-Modality (F(1,33)=8.8, p=.005) and Group-by-Language condition-by-Modality (F(1,33)=7.8, p=.009). The other interactions were not significant (Group-by-Hemisphere-by-Language condition-by-Modality F(1,33)=0.5, p=.49; Hemisphere-by-Language condition-by-Modality F(1,33)<0.01, p=.97; Group-by-Hemisphere F(1,33)=1.1, p=.31; Modality-by-Language F(1,33)=2.9, p=.10).

Most relevant to the question of group differences, the three-way Group-by-Hemisphere-by-Language condition interaction was marginal (F(1,33)=4.1, p=.05) and separate within-group ANOVAs are explored below.

*2.2 Sighted participants – within group ROI analysis*

In the sighted group, we tested responses to the language conditions in the face-preferring ROIs in FFA parcels (Hemisphere (left, right) x Language condition (words, control) x Modality (visual, audio) repeated measures ANOVA) (see Fig. 3, Fig. S1, S2 and S3). There were significant main effects of all three factors, with greater overall responses in the left rather than right hemisphere, for control stimuli over language stimuli and for visual than audio stimuli (main effect Hemisphere, F(1,14)=9.9, p=.007; main effect Language condition, F(1,14)=6.6, p=.02; main effect Modality, F(1,14)=37.4, p<.001). There was a significant Hemisphere-by-Modality interaction (F(1,14)=9.3, p=.009), but responses to visual stimuli were significant stronger than audio in both hemispheres (left, t(29)=7.8, corrected p<.001; right, t(29)=5.2, corrected p<.001).There was a significant Language condition-by-Modality interaction (F(1,14)=5.5, p=.035), with greater responses to visual false fonts than visual words (t(29)=3.6, corrected p =.002) but no difference between words and control stimuli in the audio condition (t(29)<0.1, corrected p>.99). The other interactions were not significant (Hemisphere-by-Language condition, F(1,14)<0.01, p=.9; Hemisphere-by-Language condition-by-Modality, F(1,14)=0.1, p=.75). In sum, there are no responses to language in face-preferring vertices of either hemisphere in the sighted.

We examined responses to faces in written (visual) language-preferring ROIs in FFA parcels in the sighted group with a Hemisphere (left, right) x Condition (faces, objects) repeated measures ANOVA. There was a significant main effect of Hemisphere (F(1,14)=7.2, p=.018), with stronger responses across conditions (i.e., faces and objects) in the left hemisphere. The main effect of Condition was marginal (F(1,14)=3.7, p=.08), with a trend for higher responses to control objects than faces. The Hemisphere-by-Condition interaction was not significant (F(1,14)=0.4, p=.6).

*2.2.2 Sighted responses to language conditions in visual word-preferring vOTC*

Last, we tested responses to the language conditions (Experiment 2) in visual word-preferring vertices in FFA anatomical parcels in the sighted group (top 5% visual word > false fonts, leave-one-run-out cross validation; see Fig. S1, S2 and S3). We performed a Hemisphere (left, right) x Language condition (words, control) x Modality (visual, audio) repeated measures ANOVA. There were main effects of Hemisphere (F(1,14)=8.1, p=.002) and Language condition (F(1,14)=7.3, p=.02), with stronger responses in the left than right hemisphere and overall stronger responses to visual/audio language versus control conditions (false fonts and backward speech). There was no main effect of Modality (F(1,14)=1.4, p=.3) but the Hemisphere-by-Modality interaction was significant (F(1,14)=10.0, p=.01). There were stronger responses to visual than audio stimuli in the left hemisphere (t(29)=3.0, corrected p =.01) but no difference between modalities in the right hemisphere (t(29)=0.5, corrected p >.99). The other interactions were not significant (Hemisphere-by-Language condition, F(1,14)=2.6, p=.13; Language condition-by-Modality, F(1,14)<0.1, p=.9; Hemisphere-by-Language condition-by-Modality, F(1,14)=0.1, p=.7). For completeness, post-hoc pairwise comparisons between language and control conditions for each modality: visual words vs. false fonts, t(29)=2.0, corrected p = .11; audio words vs. backward speech, t(29)=3.3, corrected p =.006.

*2.3 Blind participants – within group ROI analysis*

In the blind group, we tested responses to the language conditions in the face-preferring ROIs in FFA parcels (Hemisphere (left, right) x Language condition (words, control) x Modality (tactile, audio) repeated measures ANOVA) (see Fig. 3, Fig. S1, S2 and S3).

There were significant main effects of Hemisphere (F(1,19)=17.1, p<.001) and Language condition (F(1,19)=17.6, p<.001), reflecting stronger responses in the left than right hemisphere and stronger responses for words over control stimuli. The main effect of Modality was not significant (F(1,19)=1.7, p=.2).

There was a significant Hemisphere-by-Language condition interaction (F(1,19)=11.0, p=.004). In the left hemisphere, words elicited stronger responses than control stimuli in the face-preferring vertices (t(39)=7.0, corrected p<.001). In the right hemisphere face preferring vertices, this effect was marginal, i.e., did not survive Bonferroni correction (t(39)=2.1, corrected p=.09). The other interactions were not significant (Hemisphere-by-Modality interaction F(1,19)<0.01, p>.99), Language condition-by-Modality F(1,19)=1.3, p=.3, Hemisphere-by-Language condition-by-Modality F(1,19)=0.7, p=.4).

We also examined responses to faces in tactile language-preferring ROIs in FFA parcel in the blind group with a Hemisphere (left, right) x Condition (faces, scenes) repeated measures ANOVA. There was a significant main effect of Condition (F(1,19)=6.0, p=.02), with stronger responses to faces than tactile scenes. There was a marginal main effect of Hemisphere (F(1,19)=0.1, p=.08), suggesting stronger overall responses in the left hemisphere. There was no significant Hemisphere-by-Condition interaction (F(1,19)=0.3, p=.6).

*2.3.2 Responses to language conditions in tactile word-preferring vOTC in the blind group*

We tested responses to the language conditions (Experiment 2) in tactile word-preferring vertices in FFA anatomical parcels in the blind group (top 5% tactile word > shapes, leave-one-run-out cross validation; see Fig. S1, S2 and S3). We performed a Hemisphere (left, right) x Language condition (words, control) x Modality (tactile, audio) repeated measures ANOVA. There were main effects of Hemisphere (F(1,19)=20.5, p<.001) and Language condition (F(1,19)=64.1, p<.001), with stronger responses in the left than right hemisphere and overall stronger responses to tactile/audio language versus control conditions (tactile shapes and backward speech). There was no main effect of Modality (F(1,19)=0.1, p=.8). The Hemisphere-by-Language condition interaction was significant (F(1,19)=9.5, p=.01), but responses were stronger to words than control stimuli in both hemispheres (left, t(39)=9.0, corrected p <.001; right, t(39)=7.2, corrected p <.001). There was also a significant Language condition-by-Modality interaction (F(1,19)=38.9, p<.001), but responses to words were stronger than control conditions for both modalities (tactile, t(39)=9.2, corrected p <.001; audio, t(39)=7.1, corrected p <.001). The other interactions were not significant (Hemisphere-by-Modality, F(1,19)=0.1, p=.8; Hemisphere-by-Language condition-by-Modality, F(1,19)=2.1, p=.2). In sum, there were reliable, modality-independent language effects in both hemispheres in tactile language-preferring vOTC in blind individuals.

*3. Within-hemisphere comparisons of face- and language-effects*

*3.1 Face- and language-effects are anatomically separable in the sighted group*

In each hemisphere, in visual face-preferring ROIs in FFA parcels, we compared the face and language effects (i.e., responses to key conditions (visual faces or words) relative to their respective control conditions (objects or false fonts)) with an Experiment (1: face experiment, 2: language experiment) x Condition (key condition: faces/words, control condition: objects/false fonts) repeated measures ANOVA (Fig. S3).

There were significant Experiment-by-Condition interactions in both hemispheres (left, F(1,14)=16.1, p=.001; right, F(1,14)=20.3, p<.001). In both hemispheres, there were stronger responses to faces than objects (left, t(14)=30.8, corrected p<.001; right, t(14)=21.6, corrected p<.001) and a *reverse* language effect (preference for false fonts over words) that survived Bonferroni correction in the right hemisphere only (left, t(14)=4.9, corrected p=.09; right, t(14)=10.0, corrected p=.01). There were no significant main effects of Experiment (left, F(1,14)<0.01, p=.9; right, F(1,14)=0.3, p=.6) or Condition (left, F(1,14)=3.2, p=.1; right, F(1,14)=0.3, p=.6). Thus, in sighted people, in both hemispheres, face-preferring vOTC shows preferred responses to faces but not to language.

3.2 *Face- and language-effects are overlapping in the blind group*

In face-preferring vertices in FFA parcels in the blind group, we also compared the tactile face and language effects (i.e., responses to key conditions (tactile faces or words) relative to their respective control conditions (scenes or tactile shapes) in both hemispheres (see Fig. S3). For each hemisphere, we performed an Experiment (1: face experiment, 2: language experiment) x Condition (key condition: faces/words, control condition: scenes/shapes) repeated measures ANOVA.

Results showed the face and language effects were not different in the blind group (see Fig. S3). In both hemispheres there was a main effect of Condition (i.e., faces/words > control conditions; left, $F(1,19)=23.6$, $p<.001$; right, $F(1,19)=19.6$, $p<.001$) and no significant Experiment-by-Condition interactions (left, $F(1,19)=0.1$, $p=.8$; right, $F(1,19)=0.8$, $p=.4$). The main effects of Experiment were also not significant (left, $F(1,19)=0.3$, $p=.6$; right, $F(1,19)=0.5$, $p=.5$). Thus, in the blind group's face-preferring vOTC, face and language effects (i.e., response to key conditions relative to control conditions) were similar within both hemispheres.

*4. No difference in location of blind individuals' peak responses to faces and language in vOTC*

For the blind participants, we located the peak coordinates for responses to faces and language in vOTC within individuals using the three key contrasts (tactile faces>scenes; tactile words>shapes; audio words>backward speech). The X, Y and Z coordinates of the peaks were compared across the three contrasts using one-way repeated measures ANOVAs with Contrast (face, tactile language, spoken language) as the independent variable for each hemisphere. There were no differences across X (left, $F(2,38)=2.6$, $p=.09$; right, $F(2,38)=2.4$, $p=.1$), Y (left, $F(2,38)=0.8$, $p=.5$; right, $F(2,38)=0.1$, $p=.9$) or Z (left, $F(2,38)=0.3$, $p=.8$; right, $F(2,38)=0.1$, $p=.9$) axes. See Fig. S6 for left hemisphere peaks.

*5. Experiment 3*

*5.1 Face selectivity in an additional sighted group*

We confirmed both face-selectivity and separation of face and language responses in an additional sighted group in Experiment 3 (n=15; age (years) mean = 41.7, SD=14.2; education (years) mean = 17.2 y, SD=2.3) (see Fig. S4). Experimental procedures and analysis methods are the same as that reported in the main paper, except that the face viewing task was a dynamic face localizer from Pitcher et al. (2011). In brief, participants performed 1-back repetition detection with short video clips of faces, bodies, objects, scenes and scrambled objects. Participants also performed the language experiment described in Experiment 2.

First we tested for face-selectivity in face-preferring ROIs in the FFA anatomical parcel. We defined face-preferring ROIs using both the faces > scenes and faces > objects contrasts (top 5%, leave-one-run-out cross validation procedure) and then tested responses to the different conditions in held-out data using one-way ANOVAs (Condition: faces, scenes, objects, bodies, scrambled objects) performed for each hemisphere (Fig. S4). ANOVA results were corrected for sphericity violations where appropriate.

Face-preferring vertices defined using the faces > scenes contrast showed a main effect of Condition in both hemispheres (left, $F(2,26) = 37.5$, $p<.001$; right, $F(4,56)=48.5$, $p<.001$; post-hoc pairwise comparisons faces > scenes, bodies, objects, or scrambled objects, all $ps <.001$). The same pattern of results were seen when defining face-preferring ROIs using the faces > objects contrast (main effect of Condition: left, $F(2,23)=37.8$, $p<.001$; right, $F(4,56)=53.8$, $p<.001$, all post-hoc pairwise comparisons faces > scenes, bodies, objects, or scrambled objects, all $ps <.001$).

5.2 *Language and face responses do not overlap in sighted VOTC*

We then tested for responses to the language conditions in these face-preferring ROIs using Modality (visual, audio) by Language condition (words, control) repeated measures ANOVAs. The pattern was identical in all face-preferring ROIs in both hemispheres, regardless of contrast used to define face-selective vertices. For all face-preferring ROIs in FFA parcels, there was a main effect of Modality, reflecting stronger to visual stimuli than audio stimuli (ROIs defined faces > scenes: left, $F(1,14)=38.6$, $p<.001$; right, $F(1,14)=16.2$, $p=.001$; ROIs defined faces > objects: left, $F(1,14)=30.7$, $p<.001$; right, $F(1,14)=9.2$, $p=.01$). In all ROIs, there was no main effect of Language condition (ROIs defined faces > scenes: left, $F(1,14)=0.6$, $p=.5$; right, $F(1,14)=3.0$, $p=.1$; ROIs defined faces > objects: left, $F(1,14)=1.1$, $p=.3$; right, $F(1,14)=1.2$, $p=.3$), and no Modality-by-Language condition interaction (ROIs defined faces > scenes: left, $F(1,14)=0.3$, $p=.6$; right, $F(1,14)=1.3$, $p=.3$; ROIs defined faces > objects: left, $F(1,14)<0.01$, $p>.99$; right, $F(1,14)=0.5$, $p=.5$). Thus, in Experiment 3 visual FFAs show face-selectivity and but do not show responses to language.

Last, we confirmed a separation of responses to faces and language in Experiment 3 by identifying language-preferring ROIs in FFA anatomical parcels using the visual words > false fonts contrast (top 5%, leave-one-run-out cross validation). We tested for language responses (held-out data) in these ROIs and found preferred responses to language (i.e., language > control) only in the left hemisphere. A Modality (visual, audio) by Language condition (language, control) repeated measures ANOVA performed on visual language-preferring vertices showed main effects of Modality ($F(1,14) = 21.0$, $p<.001$) and Language condition ($F(1,14)=7.4$, $p=.02$) in the left hemisphere only (right hemisphere: marginal main effect of Modality (visual>audio), $F(1,14)=3.6$, $p=.08$), no main effect Language condition, $F(1,14)=0.5$, $p=.5$). Visual language-preferring vertices in the left hemisphere showed stronger responses to visual/audio words over the control conditions (false fonts and backward speech), and also preferred visual stimuli over audio stimuli. The interaction was also not significant in either hemisphere (left, $F(1,14)=0.1$, $p=.7$); right, $F(1,14)=2.4$, $p=.15$).

These language-preferring vertices in the left hemisphere did not prefer faces over other objects from the dynamic face localizer (one-way ANOVA with Condition: faces, scenes, objects, bodies, scrambled objects). The main effect of Condition was not significant in left hemisphere language-preferring ROIs ($F(2,25)=0.8$, $p=.4$). There was a significant main effect of Condition in the right hemisphere ROIs ($F(3,39)=3.4, p=.03$). Although these ROIs were defined as visual words > false fonts, these right hemisphere vertices did not show a preference for language compared to control conditions (see above). In the right hemisphere ROIs, there was a stronger preference for faces compared to scenes ($p=.006$) but no difference between faces and the other conditions (all $ps > .4$). In sum, language-preferring vertices in the left vOTC of sighted people prefer language and not faces, whereas in the right vOTC there are no vertices that respond to language, instead here faces are the most preferred stimulus across all ROIs tested.

**Table S1.** Demographic information for blind group.

| | Age (years) | Gender | Handed-ness | Max. education level | Education (years) | Blindness cause | Age began Braille | Braille ability (1-5) |
|---|---|---|---|---|---|---|---|---|
| 1 | 33.3 | F | Amb | Bach | 16 | LCA | 3 | 5 |
| 2 | 34.8 | F | R | PG | 19 | ROP | 3 | 3 |
| 3 | 36.9 | M | R | SC | 15 | ONH | 3 | 5 |
| 4 | 67.4 | F | R | Bach | 16 | ROP | 6 | 5 |
| 5 | 43.7 | F | R | PG | 19 | LCA | 4 | 5 |
| 6 | 57.4 | F | L | PG | 17 | LCA | 7 | 5 |
| 7 | 23.2 | M | R | Bach | 17 | CMO | 3 | 4 |
| 8 | 52.9 | M | R/Amb | Bach | 17 | Unknown, non-functional retinas | 8 | 3 |
| 9 | 32.4 | F | L | HS | 12 | LCA | 3 | 5 |
| 10 | 28.6 | F | R | Bach | 16 | ROP | 3 | 5 |
| 11 | 32.3 | F | R | PG | 20 | Congenital glaucoma | 6 | 5 |
| 12 | 49.1 | M | Amb | Bach | 17 | CMO, cornea malfunction, glaucoma | 5 | 5 |
| 13 | 33.0 | M | R/Amb | SC | 16 | LCA | 4 | 4 |
| 14 | 18.3 | M | Amb | HS | 12 | Peter's Anomaly, non-functional retinas | 3 | 3 |
| 15 | 41.3 | F | R/Amb | PG | 19 | LCA | 5 | 5 |
| 16 | 46.0 | F | R | PG | 19 | LCA | 6 | 5 |
| 17 | 50.9 | M | R | Bach | 16 | LCA | 5 | 4 |
| 18 | 46.0 | M | Amb | Bach | 17 | ROP | 3 | 5 |
| 19 | 38.8 | F | L/Amb | Bach | 16 | ONH | 5 | 5 |
| 20 | 44.7 | F | R | PG | 18 | ROP | 4 | 5 |
| Mean | 40.5 | | | | 16.7 | | 4.5 | 4.6 |
| SD | 11.8 | | | | 2.1 | | 1.5 | 0.8 |

*Note.* Age is at time of face perception experiment. Handedness: left (L), right (R), or ambidextrous (Amb) based on self-report. Abbreviations: SC = some college; Bach = bachelor degree; PG = post-graduate degree; HS = high school; ROP = retinopathy of prematurity; LCA = Leber's congenital amaurosis; ONH = optic nerve hypoplasia; CMO = congenital micro-ophthalmia. For Braille ability, participants were asked: "On a scale of 1 to 5, how well are you able to read Braille, where 1 is 'not at all', 2 is 'very little', 3 is 'reasonably well', 4 is 'proficiently', and 5 is 'expert'?"

**Table S2.** Activated clusters in whole cortex analysis in blind group.

| | | Peak MNI coordinates | | | Peak z-value | Cluster size | |
|---|---|---|---|---|---|---|---|
| | | X | Y | Z | | vertices | mm$^2$ |
| **Tactile faces > scenes** | | | | | | | |
| Left hemisphere | Postcentral gyrus | -57.1 | -15.8 | 31.2 | 5.3 | 553 | 934.3 |
| | Superior insula | -39.6 | -4.8 | 19.2 | 4.81 | | |
| | Fusiform gyrus / occipitotemporal sulcus / inferior temporal gyrus | -42.7 | -46.2 | -10.9 | 6.0 | 256 | 576.1 |
| Right hemisphere | Postcentral sulcus | 56.8 | -18.1 | 30.9 | 6.3 | 559 | 940.8 |
| | Superior insula | 34 | -9.6 | 16.8 | 5.9 | | |
| | Inferior frontal sulcus /gyrus (Parstriangularis) | 43.4 | 38.6 | 2.3 | 5.8 | 247 | 497.7 |
| **Tactile language > shapes** | | | | | | | |
| Left hemisphere | Lateral temporal / occipital / ventral occipitotemporal cortices | -41.1 | -51.4 | -16.4 | 6.2 | 3426 | 7030.5 |
| | Superior temporal sulcus / middle temporal gyrus / middle temporal sulcus | -52.2 | -35.1 | 3.6 | 4.8 | | |
| | Middle occipital gyrus | -40.8 | -76.1 | 3.6 | 4.8 | | |
| | Calcarine sulcus | -19.1 | -70.7 | 10.3 | 3.7 | | |
| | Occipitotemporal sulcus / lingual gyrus | -24.8 | -63.6 | -5.1 | 4.8 | | |
| | Fusiform gyrus / lateral occipitotemporal sulcus | -41.1 | -51.4 | -16.4 | 6.2 | | |

| | | | | | | | |
|---|---|---|---|---|---|---|---|
| | Inferior frontal gyrus (Parstriangularis) | -48.7 | 28.2 | 6.3 | 4.7 | 720 | 1452.9 |
| | Inferior frontal sulcus / inferior precentral sulcus | -33.5 | 5.8 | 28.2 | 3.8 | | |
| Right hemisphere | Superior temporal sulcus, middle temporal gyrus | 38.2 | -60.1 | 16.3 | 6.1 | 439 | 787.6 |
| | Fusiform gyrus, occipitotemporal gyrus / sulcus | 41.1 | -48.2 | -15.9 | 4.2 | 235 | 516.9 |
| **Spoken language > backward speech** | | | | | | | |
| Left hemisphere | Lateral temporal / occipital / ventral occipitotemporal cortices | -48.5 | -38.5 | 4.5 | 6.1 | 2732 | 5520.1 |
| | Superior temporal sulcus / gyrus, middle temporal gyrus | -48.5 | -38.5 | 4.5 | 6.1 | | |
| | Anterior occipital sulcus | -39.5 | -71.5 | 6.6 | 5.1 | | |
| | Inferior temporal gyrus / fusiform gyrus / lateral occipitotemporal sulcus | -41.1 | -50 | -14.4 | 6.0 | | |
| | Inferior frontal gyrus / sulcus (Parstriangularis) | -51.9 | 29.9 | 5.5 | 5.2 | 441 | 883.4 |
| | Inferior precentral sulcus | -38.2 | 4.2 | 24.8 | 3.6 | | |
| Right hemisphere | Superior temporal sulcus, middle temporal gyrus | 55.9 | -53.4 | 9.6 | 5.3 | 815 | 1529.0 |
| | Superior temporal sulcus | 47.5 | -29.4 | 2.5 | 5.0 | | |

| | | | | | | | |
|---|---|---|---|---|---|---|---|
| | Calcarine sulcus, lingual gyrus | 15.9 | -74.5 | 8.2 | 4.2 | 245 | 701.8 |
| | Fusiform gyrus, occipitotemporal gyrus / sulcus, transverse collateral sulcus | 41.0 | -45.3 | -18.1 | 4.0 | 353 | 764.6 |

*Note*. Coordinates are in MNI152 space.
